# Personalized deep learning of individual immunopeptidomes to identify neoantigens for cancer vaccines

**DOI:** 10.1101/620468

**Authors:** Ngoc Hieu Tran, Rui Qiao, Lei Xin, Xin Chen, Baozhen Shan, Ming Li

## Abstract

Tumor-specific neoantigens play the main role for developing personal vaccines in cancer immunotherapy. We propose, for the first time, a personalized *de novo* sequencing workflow to identify HLA-I and HLA-II neoantigens directly and solely from mass spectrometry data. Our workflow trains a personal deep learning model on the immunopeptidome of an individual patient and then uses it to predict mutated neoantigens of that patient. This personalized learning and mass spectrometry-based approach enables comprehensive and accurate identification of neoantigens. We applied the workflow to datasets of five melanoma patients and substantially improved the accuracy and identification rate of *de novo* HLA peptides by 14.3% and 38.9%, respectively. This subsequently led to the identification of 10,440 HLA-I and 1,585 HLA-II new peptides that were not presented in existing databases. Most importantly, our workflow successfully discovered 17 neoantigens of both HLA-I and HLA-II, including those with validated T cell responses and those novel neoantigens that had not been reported in previous studies.

## INTRODUCTION

Neoantigens are tumor-specific mutated peptides that are brought to the surface of tumor cells by major histocompatibility complex (MHC) proteins and can be recognized by T cells as “foreign” (non-self) to trigger immune response. As neoantigens carry tumor-specific mutations and are not found in normal tissues, they represent ideal targets for the immune system to distinguish cancer cells from non-cancer ones [1-3]. The potential of neoantigens for cancer vaccines is supported by multiple evidences, including the correlation between mutation load and response to immune checkpoint inhibitor therapies [4, 5], neoantigen-specific T cell responses detected even before vaccination (naturally occurring) [6-8]. Indeed, three independent studies have further demonstrated successful clinical trials of personalized neoantigen vaccines for patients with melanoma [6-8]. The vaccination was found to reinforce pre-existing T cell responses and to induce new T cell populations directed at the neoantigens. In addition to developing cancer vaccines, neoantigens may help to identify targets for adoptive T cell therapies, or to improve the prediction of response to immune checkpoint inhibitor therapies.

And thus began the “gold rush” for neoantigen mining [1-3]. The current prevalent approach to identify candidate neoantigens often includes two major phases: (i) exome sequencing of cancer and normal tissues to find somatic mutations and (ii) predicting which mutated peptides are most likely to be presented by MHC proteins for T cell recognition. The first phase is strongly backed by high-throughput sequencing technologies and bioinformatics pipelines that have been well established through several genome sequencing projects during the past decade. The second phase, however, is still facing challenges due to our lack of knowledge of the MHC antigen processing pathway: how mutated proteins are processed into peptides; how those peptides are delivered to the endoplasmic reticulum by the transporter associated with antigen processing; and how they bind to MHC proteins. To make it further complicated, human leukocyte antigens (HLA), those genes that encode MHC proteins, are located among the most genetically variable regions and their alleles basically change from one individual to another. The problem is especially more challenging for HLA class II (HLA-II) peptides than HLA class I (HLA-I), because the former are longer, their motifs have greater variations, and very limited data is available.

Current *in silico* methods focus on predicting which peptides bind to MHC proteins given the HLA alleles of a patient, e.g. NetMHC [9, 10]. However, usually very few, less than a dozen from thousands of predicted candidates are confirmed to be presented on the tumor cell surface and even less are found to trigger T cell responses, not to mention that real neoantigens may not be among top predicted candidates [1, 2]. Several efforts have been made to improve the MHC binding prediction, including using mass spectrometry data in addition to binding affinity data for more accurate prediction of MHC antigen presentation [11, 12, 26-28]. Recently, proteogenomic approaches have been proposed to combine mass spectrometry and exome sequencing to identify neoantigens directly isolated from MHC proteins, thus overcoming the limitations of MHC binding prediction [13, 14]. In those approaches, exome sequencing was performed to build a customized protein database that included all normal and mutated protein sequences. The database was further used by a search engine to identify endogenous peptides, including neoantigens, that were obtained by immunoprecipitation assays and mass spectrometry.

Existing database search engines, however, are not designed for HLA peptides and may be biased towards tryptic peptides [15, 16]. They may have sensitivity and specificity issues when dealing with a very large search space created by (i) all mRNA isoforms from exome sequencing and (ii) unknown digestion rules for HLA peptides. Furthermore, recent proteogenomic studies reported a weak correlation between proteome- and genome-level mutations, where the number of identified mutated HLA peptides was three orders of magnitudes less than the number of somatic mutations that were provided to the database search engines [13,14]. A large number of genome-level mutations were not presented at the proteome level, while at the same time, some mutated peptides might be difficult to detect at the genome level. For instance, Faridi *et al.* found evidence of up to 30% of HLA-I peptides that were cis- and trans-splicing, which couldn’t be detected by exome sequencing nor protein database search [25]. Thus, an independent approach that does not rely heavily on genome-level information to identify mutated peptides directly from mass spectrometry data is needed, and *de novo* sequencing is the key to address this problem.

In this study, we propose, for the first time, a personalized *de novo* sequencing workflow to identify HLA-I and HLA-II neoantigens directly and solely from mass spectrometry data. *De novo* sequencing is invented for the purpose of discovering novel peptides and proteins, genetic variants or mutations. Thus, its application to identify neoantigens is a perfect match. We bring *de novo* sequencing to the “personalized” level by training a specific machine learning model for each individual patient using his/her own data. In particular, we use the collection of normal HLA peptides, i.e. the immunopeptidome, of a patient to train a model and then use it to predict mutated HLA peptides of that patient. Learning an individual’s immunopeptidome is made possible by our recent deep learning model, DeepNovo [17, 18], which uses a long-short-term memory (LSTM) recurrent neural network (RNN) to capture sequence patterns in peptides or proteins, in a similar way to natural languages [19]. This personalized learning workflow significantly improves the accuracy of *de novo* sequencing for comprehensive and reliable identification of neoantigens. Furthermore, our *de novo* sequencing approach predicts peptides solely from mass spectrometry data and does not depend on genomic information as existing approaches. We applied the workflow to datasets of five melanoma patients and successfully identified in average 154 HLA-I and 47 HLA-II candidate neoantigens per patient, including those with validated T cell responses and those novel neoantigens that had not been reported in previous studies.

## RESULTS

### Personalized *De novo* Sequencing of Individual Immunopeptidomes

Figure 1 describes five steps of our personalized *de novo* sequencing workflow to predict HLA peptides of an individual patient from mass spectrometry data: (1) build the immunopeptidome of the patient; (2) train personalized machine learning model; (3) personalized *de novo* sequencing; (4) quality control of *de novo* peptides; and (5) neoantigen selection. The details of each step on two example datasets, HLA-I and HLA-II of patient Mel-15 from [13], are provided in Supplementary Table S1 for illustration.

**Figure 1.**
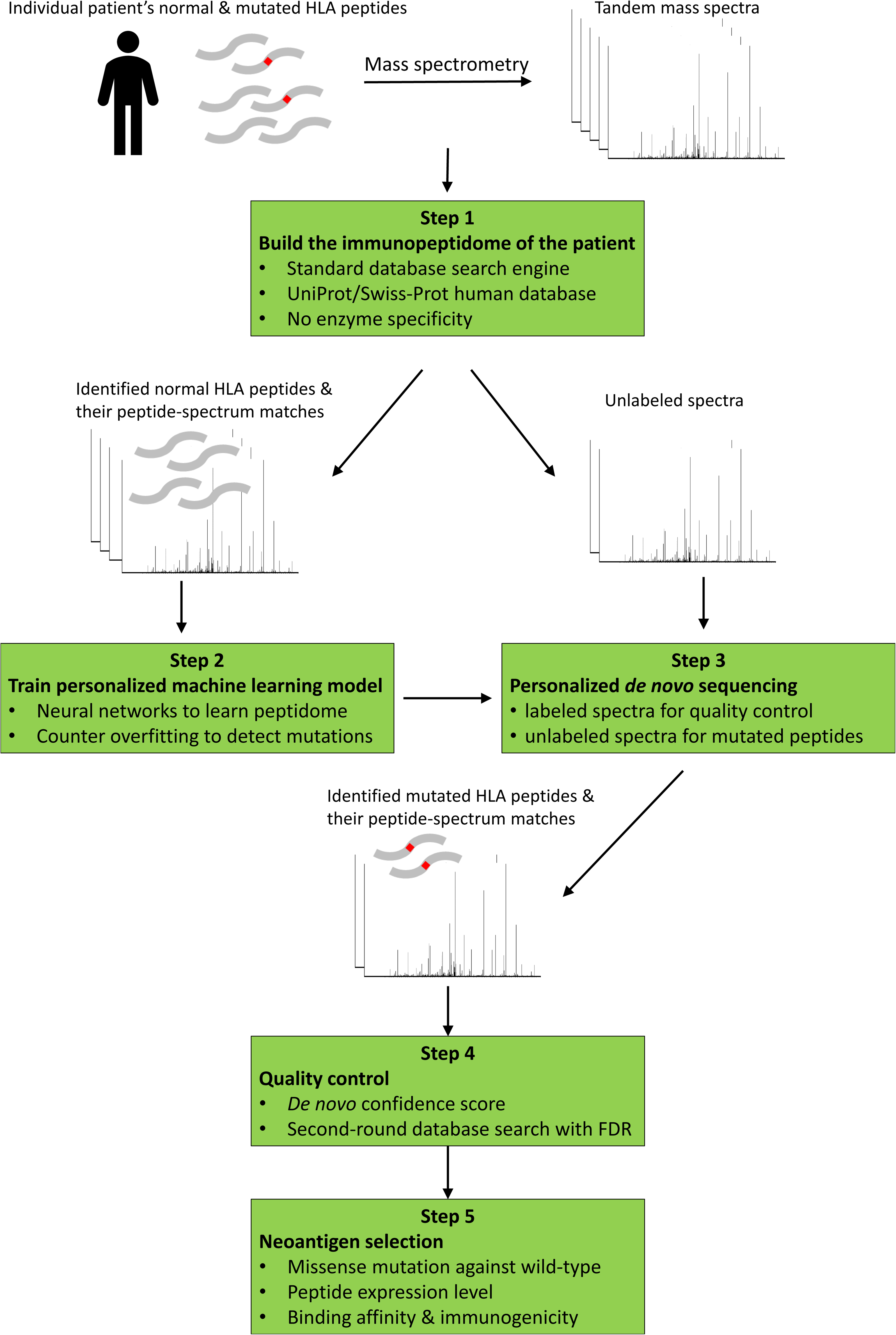
Personalized *de novo* sequencing workflow for neoantigen discovery. (HLA: Human Leukocyte Antigen; FDR: False Discovery Rate).

In step 1 of the workflow, to build the immunopeptidome of the patient, we searched the mass spectrometry data against the standard Swiss-Prot human protein database. As digestion rules for HLA peptides are unknown, a search engine that supports no-enzyme-specific digestion is needed (we used PEAKS X [18]). Identified normal HLA peptides and their peptide-spectrum matches (PSMs) at 1% false discovery rate (FDR) represent the patient’s immunopeptidome and its spectral library. Mutated HLA peptides were not presented in the protein database, so their spectra remained unlabeled. For example, we identified 341,216 PSMs of 35,551 HLA-I peptides and 67,021 PSMs of 9,664 HLA-II peptides from Mel-15 datasets. The numbers of unlabeled spectra were 596,915 and 135,490, respectively (Supplementary Table S1).

In step 2, we used the identified normal HLA peptides and their PSMs to train DeepNovo, a neural network model for *de novo* peptide sequencing [17, 18]. In addition to capturing fragment ions in tandem mass spectra, DeepNovo learns sequence patterns of peptides by modelling them as a special language with an alphabet of 20 amino acid letters. This unique advantage allowed us to train a personalized model to adapt to a specific immunopeptidome of an individual patient and achieved much better accuracy than a generic model (results are shown in a later section). At the same time, it was essential to apply counter-overfitting techniques so that the model could predict new peptides that it had not seen during training. We partitioned the PSMs into training, validation, and test sets (ratio 90-5-5, respectively) and restricted them not to share common peptide sequences. We stopped the training process if there was no improvement on the validation set and evaluated the model performance on the test set. As a result, our personalized model was able to both achieve very high accuracy on an individual immunopeptidome and detect mutated peptides. This approach is particularly useful for missense mutations (the most common source of neoantigens) because they still preserve most patterns in the peptide sequences.

In step 3, we used the personalized DeepNovo model to perform *de novo* peptide sequencing on both labeled spectra (i.e., the PSMs identified in step 1) and unlabeled spectra. Results from labeled spectra were needed for accuracy evaluation and calibrating prediction confidence scores. Peptides identified from unlabeled spectra ***<u>and</u>*** not presented in the protein database were defined as “*de novo* peptides” and would be further analyzed in the next steps to find candidate neoantigens of interest.

In step 4, a quality control procedure was designed to select high-confidence *de novo* peptides and to estimate their FDR. We first calculated the accuracy of *de novo* sequencing on the test set of PSMs by comparing the predicted peptide to the true one for each spectrum. DeepNovo also provides a confidence score for each predicted peptide, which can be used as a filter for better accuracy. Since the test set did not share common peptides with the training set, we expected the distribution of accuracy versus confidence score on the test set to be close to that of *de novo* peptides which the model had not seen during training. Thus, we calculated a score threshold at a precision of 95% on the test set and used it to select high-confidence *de novo* peptides (Figure 2b). Finally, to estimate the FDR of high-confidence *de novo* peptides, we performed a second-round PEAKS X search of all spectra against a combined list of those peptides and the database peptides (i.e. normal HLA peptides identified in step 1). Only *de novo* peptides identified at 1% FDR were retained. Thus, our stringent procedure of quality control guaranteed that each *de novo* peptide was supported by solid evidences from two independent tools, DeepNovo and PEAKS X. For instance, we found 16,226 HLA-I and 2,717 HLA-II high-confidence *de novo* peptides from Mel-15 datasets. Among them, 5,320 HLA-I and 863 HLA-II *de novo* peptides passed 1% FDR filter (Supplementary Table S1).

**Figure 2.**
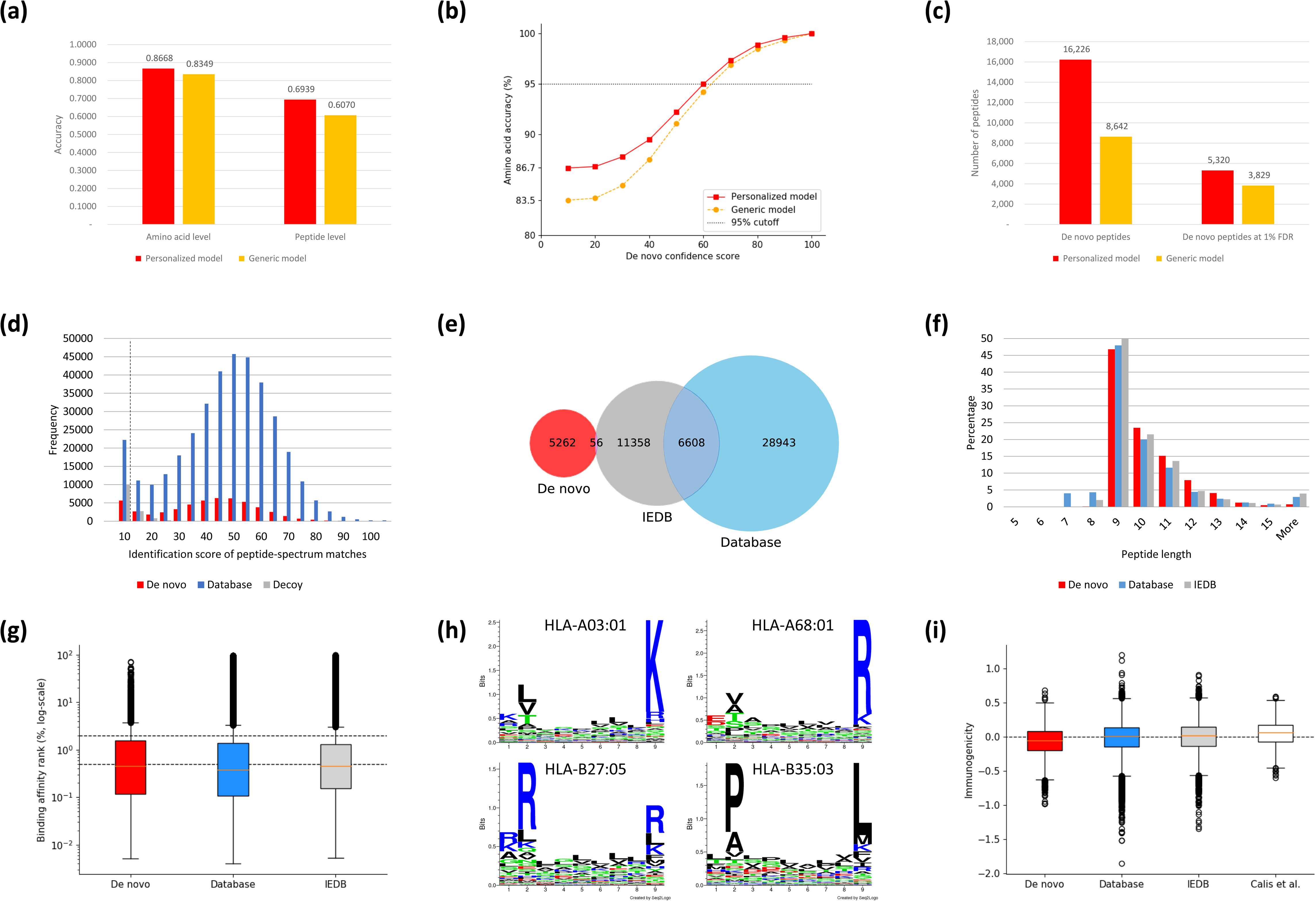
Accuracy and immune characteristics of *de novo* HLA-I peptides from patient Mel-15 dataset. (a) Accuracy of *de novo* peptides predicted by personalized and generic models. (b) Distribution of amino acid accuracy versus DeepNovo confidence score for personalized and generic models. (c) Number of *de novo* peptides identified at high-confidence threshold and at 1% FDR by personalized and generic models. (d) Distribution of identification scores of *de novo*, database, and decoy peptide-spectrum matches. The dashed line indicates 1% FDR threshold. (e) Venn diagram of *de novo*, database, and IEDB peptides. (f) Length distribution of *de novo*, database and IEDB peptides. (g) Distribution of binding affinity ranks of *de novo*, database, and IEDB peptides. Lower rank indicates better binding affinity. The two dashed lines correspond to the ranks of 0.5% and 2%, which indicate strong and weak binding, respectively, by NetMHCpan. (h) Binding sequence motifs identified from *de novo* peptides by GibbsCluster. (i) Immunogenicity distribution of *de novo*, database, IEDB, and Calis *et al.*’s peptides [24]. (HLA: Human Leukocyte Antigen; FDR: False Discovery Rate; IEDB: Immune Epitope Database).

We applied this workflow to train personalized models for another four patients and to predict their HLA-I and HLA-II *de novo* peptides. The five patients, Mel-5, Mel-8, Mel-12, Mel-15, Mel-16, were selected because their neoantigens had been identified and validated by a proteogenomic database search approach in [13]. In total, we identified 10,440 HLA-I and 1,585 HLA-II *de novo* peptides at 1% FDR (Table 1). The number of identified database peptides in step 1 of the workflow was 97,526 HLA-I and 15,835 HLA-II. Thus, our *de novo* sequencing results expanded the immunopeptidomes by approximately 10%.

**Table 1.** Number of *de novo* and database HLA peptides identified at 1% FDR.

| Patient ID | HLA-I |  | HLA-II |  |
| --- | --- | --- | --- | --- |
|  | Database | *De novo* | Database | *De novo* |
| Mel-5 | 12,998 | 1,272 | MS data not available |  |
| Mel-8 | 13,635 | 1,235 | MS data not available |  |
| Mel-12 | 10,068 | 1,354 | MS data not available |  |
| Mel-15 | 35,551 | 5,320 | 9,664 | 863 |
| Mel-16 | 25,274 | 1,259 | 6,171 | 722 |

FDR: False Discovery Rate
MS: Mass Spectrometry

### Advantages of Personalized Model over Generic Model

To demonstrate the advantages of our personalized approach, we compared the personalized model of patient Mel-15’s HLA-I to a generic model, which had the same neural network architecture but was trained on a combined HLA-I dataset of 9 other patients from the same study [13]. All datasets were derived from the same experiment and instrument, the only difference is the immunopeptidomes of the patients. The combined dataset has 477,482 PSMs, which is 39.9% larger than the Mel-15 dataset (477,482 / 341,216 = 1.399). Figure 1a shows the accuracy of the personalized model versus the generic model on the Mel-15 test set. As mentioned earlier, this test set did not share common peptides with the Mel-15 training set, so both models had not seen the test peptides during training. The personalized model achieved 14.3% higher accuracy at the peptide level (0.6939 / 0.6070 = 1.143) and 3.8% higher accuracy at the amino acid level (0.8668 / 0.8349 = 1.038), despite its smaller training set. The superiority of the personalized model over the generic one can also be seen from the accuracy-versus-score distribution in Figure 2b. At the same level of amino acid accuracy, e.g. 95%, the personalized model required a lower score cutoff, thus allowing more *de novo* peptides to be identified. Indeed, Figure 1c shows that the personalized model identified 87.8% more high-confidence *de novo* peptides (16,226 / 8,642 = 1.878) and 38.9% more *de novo* peptides at 1% FDR (5,320 / 3,829 = 1.389). More importantly, the personalized model was able to capture 6 of 8 target neoantigens of patient Mel-15 (Table 2), while the generic model only recovered 3 of them. Those results demonstrate that our personalized approach substantially improves the accuracy and identification rate of *de novo* peptides by adapting to a specific immunopeptidome of an individual patient.

**Table 2.**
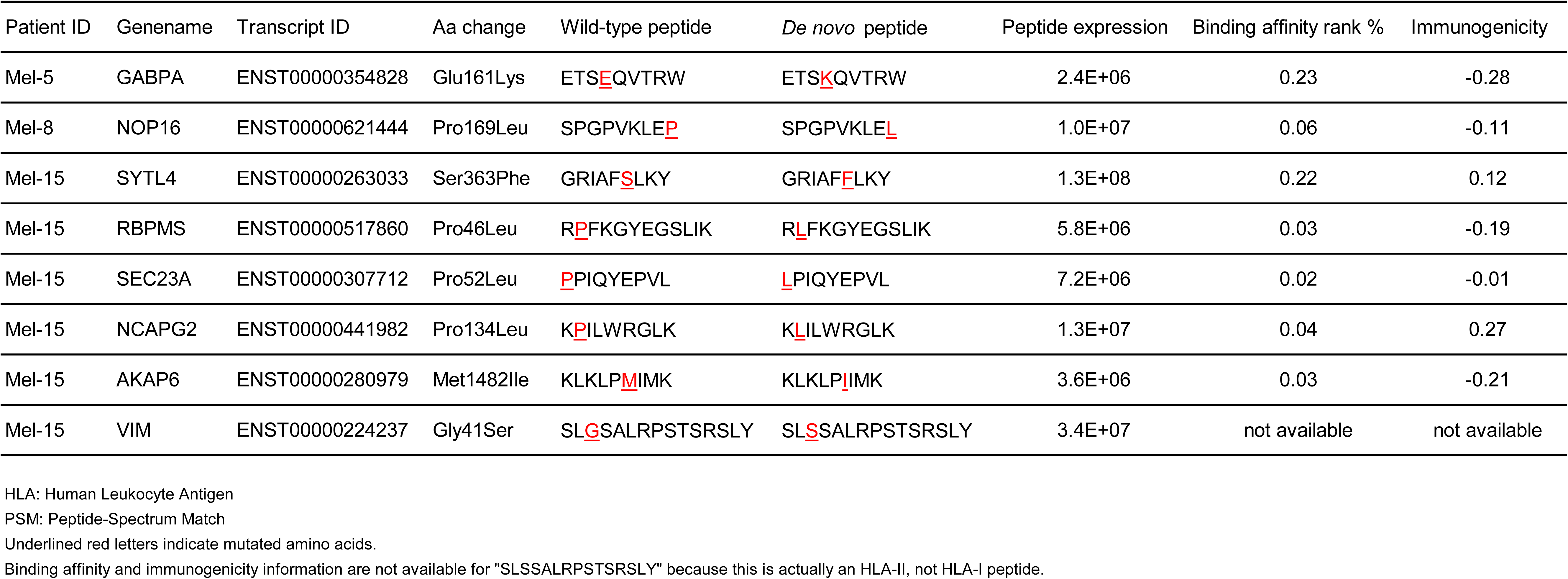
HLA-I candidate neoantigens that matched to RNA-Seq results.

It should be noted that the performance of the generic model could be improved if it is trained on a more diverse, pan-allelic dataset rather than on a limited set of 9 patients. However, such a dataset to account for large variations of HLA alleles and derived from the same mass spectrometry experiments does not exist to date. We also envision that a personalized model could be trained on top of a well pre-trained generic model to achieve better performance, reduce training time, and especially when limited data is available for an individual patient.

### Analysis of Immune Characteristics of *De novo* HLA peptides

In this section, we studied common immune features of *de novo* HLA peptides and compared them to normal HLA peptides, i.e. those identified by the database search engine in step 1 of the workflow. We also compared to previously reported human epitopes from the Immune Epitope Database (IEDB) [20].

Figure 2d shows the distribution of PEAKS X identification scores of *de novo* PSMs against those of database and decoy PSMs for HLA-I peptides of patient Mel-15. The distributions confirm that the *de novo* peptides have strong supporting PSMs as the database peptides and are clearly distinguishable from the decoy ones. The supporting PSMs of all *de novo* HLA peptides of five patients are provided in Supplementary Table S2.

Next, we compared *de novo* and database HLA-I peptides of patient Mel-15 to 18,022 IEDB epitopes, which were retrieved according to the patient’s six alleles (HLA-A03:01, HLA-A68:01, HLA-B27:05, HLA-B35:03, HLA-C02:02, HLA-C04:01). The Venn diagram in Figure 2e shows that 56 *de novo* peptides have been reported as epitopes in earlier studies. Note that the *de novo* peptides were specific to an individual patient and were not presented in the protein database, so the chance to find them in IEDB is rare. Even 81.4% (28,943 / 35,551) of the database peptides were not found in IEDB. This is due to the large variation of HLA peptides and further emphasizes the importance of our personalized approach. Figure 2e further shows that both *de novo* and database peptides have the same characteristic length distribution as IEDB epitopes. For the other four patients Mel-5, Mel-8, Mel-12, and Mel-16, we also found that the length distributions of their *de novo* HLA-I peptides are very similar to those of database peptides (Supplementary Figures S1a-d). However, for HLA-II, the *de novo* peptides tend to be longer than the database ones (Supplementary Figures S1e, f). We hypothesize that it might be challenging for the database search engine to identify long HLA-II peptides when the digestion rule is unknown.

One of the most widely used measures to assess HLA peptides is their binding affinity to MHC proteins. We used NetMHCpan [10] to predict the binding affinity of the *de novo*, database, and IEDB peptides for HLA-I alleles of patient Mel-15. Figures 2g shows that the *de novo* peptides have the same level of binding affinity as database and IEDB peptides (p-value > 0.23 for Mann-Whitney U test between *de novo* and IEDB peptides). Furthermore, majority of the *de novo* peptides were predicted as good binders by multiple criteria: 79.3% (4,220 / 5,320) weak-binding, 51.8% (2,757 / 5,320) strong-binding, and 74.0% (3,938 / 5,320) with binding affinity less than 500 nM (Supplementary Figure S2). Similar results were observed for *de novo* peptides of different HLA-I alleles of the other four patients (Supplementary Figure S3). We also applied GibbsCluster [23], an unsupervised alignment and clustering method to identify binding motifs without the need of HLA allele information. We found that the *de novo* peptides of patient Mel-15 were clustered into four groups of which motifs corresponded exactly to four alleles of the patient (Figure 2h). Note that both *de novo* sequencing and unsupervised clustering methods do not use any prior knowledge such as protein database or HLA allele information, yet their combination still revealed the correct binding motifs of the patient. This suggests that our workflow can be used to identify novel HLA peptides of unknown alleles. Results from the database peptides also yielded the same binding motifs (Supplementary Figure S4).

Finally, we used an IEDB tool (http://tools.iedb.org/immunogenicity/) [24] to predict the immunogenicity of *de novo* HLA-I peptides and then compared to database, IEDB, and human immunogenic peptides that were used in that original study (Figure 2i). We found that 38.8% (2,065 / 5,320) of the *de novo* peptides had positive predicted immunogenicity (log-likelihood ratio of immunogenic over non-immunogenic [24]). The *de novo* peptides had lower predicted immunogenicity than the database and IEDB peptides, which in turn were less immunogenic than the original peptides (Calis *et al.*). This was expected because the tool had been developed on a limited set of a few thousands well-studied peptides. The predicted immunogenicity of *de novo* HLA-I peptides of the other four patients are provided in Supplementary Figure S5.

Overall, our analysis results confirmed the correctness, and more importantly, the essential characteristics of *de novo* HLA peptides for immunotherapy. The remaining question is to select candidate neoantigens from *de novo* HLA peptides based on their characteristics.

### Neoantigen Selection and Evaluation

We considered several criteria that had been widely used in previous studies for neoantigen selection [6-8, 13, 14, 21, 22]. Specifically, we checked whether a *de novo* HLA peptide carried one amino acid substitution by aligning its sequence to the Swiss-Prot human protein database, and whether that substitution was caused by one single nucleotide difference in the encoding codon. In this paper, we refer to those substitutions as “missense-like mutations”. For each mutation, we recorded whether the wild-type peptide was also detected and whether the mutated amino acid was located at a flanking position. For expression level information of a peptide, we calculated the number of its PSMs, their total identification score, and their total abundance. Finally, we used NetMHCpan and IEDB tools [10, 24] to predict the binding affinity and the immunogenicity of a peptide. The results for 10,440 HLA-I and 1,585 HLA-II *de novo* peptides of five patients are provided in Supplementary Table S3.

To select candidate neoantigens, we focused on *de novo* HLA peptides that carried one single missense-like mutation. This criterion reduced the number of peptides considerably, e.g. from 5,320 to 328 HLA-I and from 863 to 154 HLA-II peptides of patient Mel-15. We further filtered out peptides with only one supporting PSM or with mutations at flanking positions because they were more error-prone and less stable to be effective neoantigens. In average, we obtained 154 HLA-I and 47 HLA-II candidates per patient. Expression level, binding affinity, and immunogenicity can be further used to prioritize candidates for experimental validation of immune response; we avoided using those information as hard filters (Supplementary Table S3).

We cross-checked our *de novo* HLA peptides against the nucleotide mutations and mRNA transcripts in the original publication [13]. We identified seven HLA-I and ten HLA-II candidate neoantigens that matched missense variants detected from exome sequencing (Tables 2 and 3). The first seven were among eleven neoantigens reported by the authors using a proteogenomic approach that required both exome sequencing and proteomics database search. Two HLA-I neoantigens, “GRIAF**<u>F</u>**LKY” and “K**<u>L</u>**ILWRGLK”, had been experimentally validated to elicit specific T-cell responses. We indeed observed that those two peptides had superior immunogenicity, and especially, expression level of up to one order of magnitude higher than the other neoantigens (Table 2). This observation confirms the critical role of peptide-level expression for effective immunotherapy, in addition to immunogenicity and binding affinity.

**Table 3.** HLA-II candidate neoantigens that matched to RNA-Seq results.

| ...S Y V T T S T R T Y S L <u>G</u> S A L R P S T S R S L Y... | Peptide expression | Binding affinity rank % |
| --- | --- | --- |
| Y V T T S T R T Y S L <u>S</u> S A L R P S T S | 4.3E+06 | 1.8 |
| V T T S T R T Y S L <u>S</u> S A L R P S T S | 2.3E+06 | 1.8 |
| S L <u>S</u> S A L R P S T S R S L Y | 1.7E+07 | 4.5 |
| S Y V T T S T R T Y S L <u>S</u> S A L R P S T S | 9.8E+06 | 1.7 |
| V T T S T R T Y S L <u>S</u> S A L R P S | 3.0E+06 | 1.7 |
| T T S T R T Y S L <u>S</u> S A L R P S | 1.1E+07 | 1.0 |
| Y V T T S T R T Y S L <u>S</u> S A L R P S T | 3.7E+05 | 2.0 |
| T T S T R T Y S L <u>S</u> S A L R P S T S | 7.5E+05 | 1.6 |
| Y V T T S T R T Y S L <u>S</u> S A L R P S | 8.5E+05 | 2.5 |
| T S T R T Y S L <u>S</u> S A L R P S | 1.3E+07 | 0.6 |

HLA: Human Leukocyte Antigen The mutated amino acid is underlined and highlighted in red for the reference (top row) and the neoantigens. This mutation site is located at Gly41Ser, transcript ENST00000224237, gene name VIM, patient Mel-15.

The ten HLA-II candidate neoantigens were novel and had not been reported in [13]. They were clustered around a single missense mutation and were a good example to illustrate the complicated digestion of HLA-II peptides (Table 3). Eight of them were predicted as strong binders by NetMHCIIpan (rank <= 2%), two as weak binders (rank <= 10%). The peptide located at the center of the cluster, “TSTRTYSL**<u>S</u>** SALRPS”, showed both highest expression level and binding affinity, thus representing a promising target for further experimental validation. Interestingly, another peptide, “SL**<u>S</u>**SALRPSTSRSLY”, showed up in both HLA-I and HLA-II datasets with very high expression level (Tables 2 and 3). Using a consensus method of multiple binding prediction tools from IEDB to double-check, we found that this peptide had a binding affinity rank of 0.08%, instead of 4.5% as predicted by NetMHCIIpan, and exhibited a different binding motif from the rest of the cluster. Thus, given its superior binding affinity and expression level, this peptide would also represent a great candidate for immune response validation.

We also investigated the four HLA-I neoantigens that had been reported in [13] but were not detected by our method. Three of them were not supported by good PSMs, and in fact, DeepNovo and PEAKS X identified alternative peptides that better matched the corresponding spectra (Supplementary Figure S6). The remaining neoantigen was missed due to a *de novo* sequencing error. We noticed that all four peptides had been originally identified at 5% FDR instead of 1%, so their signals were possibly too weak for identification.

## DISCUSSION

In this study, we proposed a personalized *de novo* sequencing workflow to identify HLA neoantigens directly and solely from mass spectrometry data. The key advantage of our method is the ability of its deep learning model to adapt to a specific immunopeptidome of an individual patient. This personalized approach greatly improved the performance of *de novo* sequencing and allowed to accurately identify mutated HLA peptides. For instance, we showed that a personalized model achieved up to 14.3% higher accuracy than a generic model, identified 38.9% more *de novo* peptides at 1% FDR, and doubled the number of validated neoantigens. We applied the workflow to five melanoma patients and identified 10,440 HLA-I and 1,585 HLA-II *de novo* peptides at 1% FDR, expanding their immunopeptidomes by approximately 10%. Our analysis also demonstrated that the *de novo* HLA peptides exhibited the same immune characteristics as previously reported human epitopes, including binding affinity, immunogenicity, and expression level, which are essential for effective immunotherapy. We cross-checked our *de novo* HLA peptides against exome sequencing results and found ten novel HLA-II neoantigens that had not been reported earlier. This result demonstrated the capability of our *de novo* sequencing approach to overcome the challenges of unknown digestion rules and binding prediction for HLA-II peptides. Last but not least, our *de novo* sequencing workflow directly predicted neoantigens from mass spectrometry data and did not require genome-level information nor HLA alleles of the patient as in existing approaches. Such an independent approach allows to discover novel mutated peptides that might be difficult to detect at the genome level, e.g. cis- and trans-spliced peptides. Thus, our personalized *de novo* sequencing workflow to predict mutated peptides from the cancer cell surface presents a simple and direct solution to discover neoantigens for cancer immunotherapy. As Newton said, “nature is pleased with simplicity”.

## Acknowledgements

This work was funded in part by the NSERC OGP0046506 grant and the Canada Research Chair program. N.H.T. was supported by the Mitacs Elevate Fellowship. The authors thank Dr. Kwok Pui Choi for proofreading the manuscript.

## Author Contributions

M.L. and B.S. conceived the research idea. N.H.T. designed the neoantigen discovery workflow. N.H.T and R.Q. implemented the software and analyzed the results. X.C. and L.X. contributed to model design, software development, and data analysis. N.H.T, M.L., and R.Q. wrote the manuscript. M.L., B.S., and L.X. supervised the research project.

## Software Availability

DeepNovo and the workflow are implemented in Python. The latest version is open-source and available on GitHub (https://github.com/nh2tran/DeepNovoAA).

