## Supplementary Figures and Tables for "Personalized deep learning of individual immunopeptidomes to identify neoantigens for cancer vaccines": Figure_S1.pdf

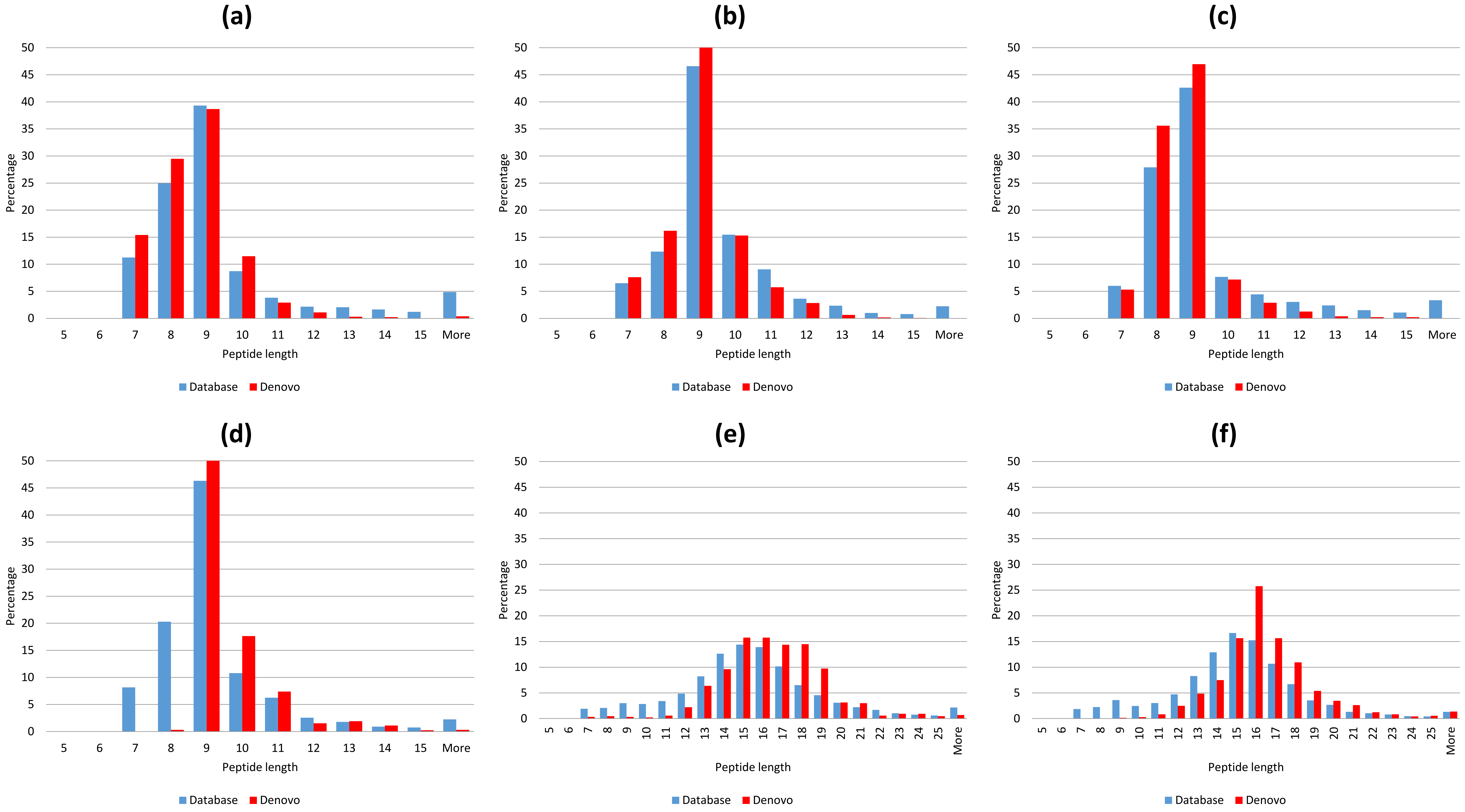


Supplementary Figure S1. Length distributions of HLA *de novo* and database peptides. (a) Mel-5 HLA-I; (b) Mel-8 HLA-I; (c) Mel-12 HLA-I; (d) Mel-16 HLA-I; (e) Mel-15 HLA-II; (f) Mel-16 HLA-II.
