## Supplementary Figures and Tables for "Personalized deep learning of individual immunopeptidomes to identify neoantigens for cancer vaccines": Figure_S2.pdf

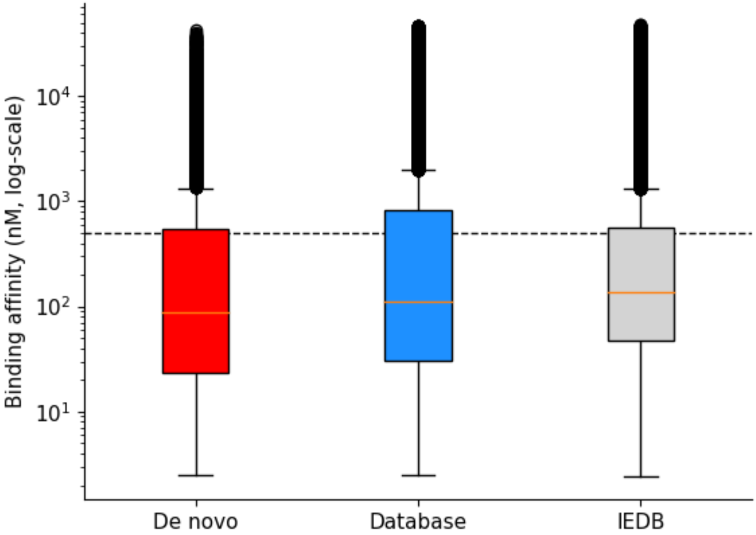


Supplementary Figure S2. Binding affinity distributions of *de novo*, database, and IEDB HLA-I peptides of patient Mel-15. The dashed line indicates the value of 500 nM, a common threshold to select good binders.
