## Supplementary Figures and Tables for "Personalized deep learning of individual immunopeptidomes to identify neoantigens for cancer vaccines": Figure_S3.pdf

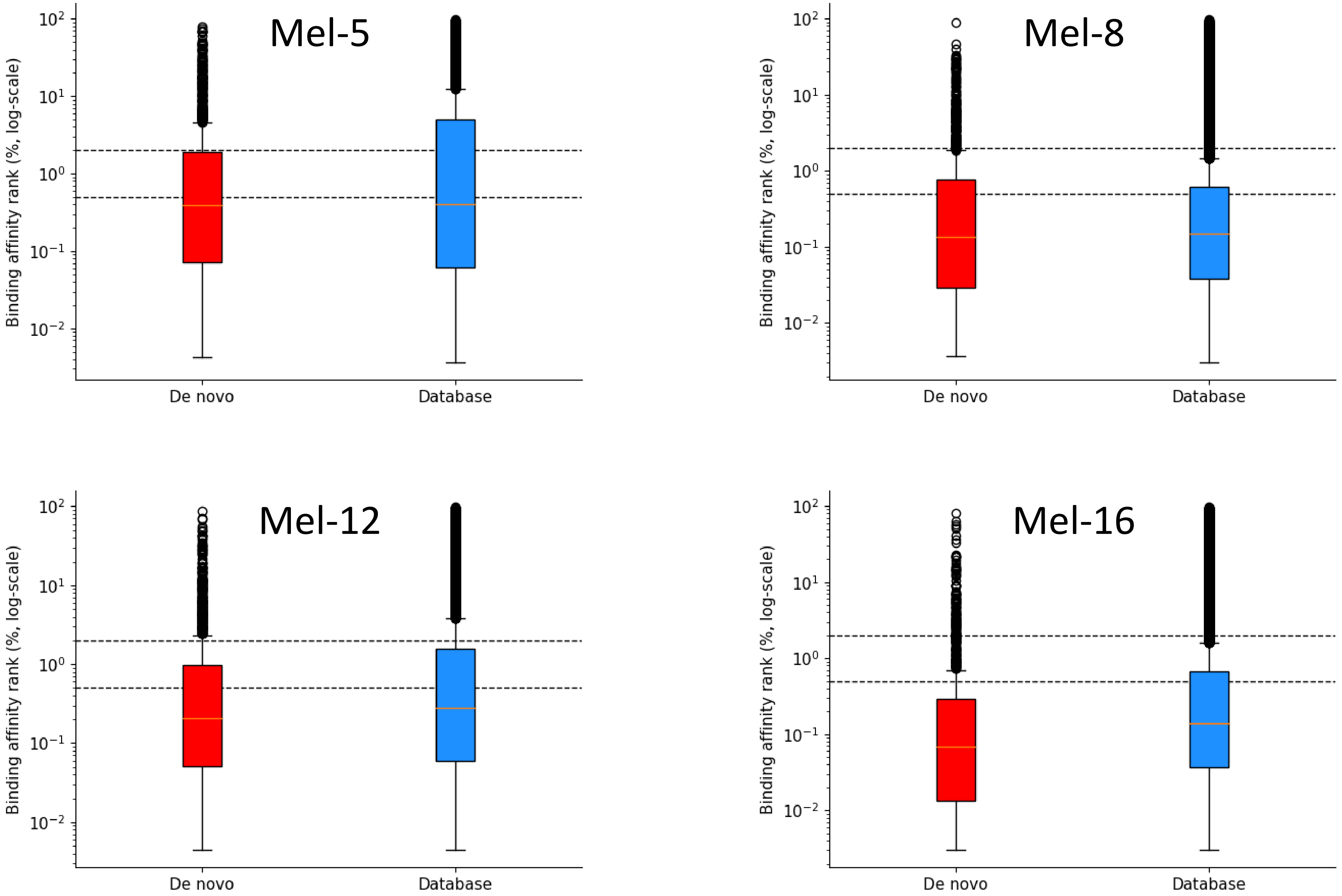


Supplementary Figure S3. Binding affinity of *de novo* and database HLA-I peptides. Dashed lines indicate default thresholds of weak-binding (rank 2.0%) and strong-binding (rank 0.5%) of NetMHCpan.
