## Supplementary Figures and Tables for "Personalized deep learning of individual immunopeptidomes to identify neoantigens for cancer vaccines": Figure_S6.pdf

Supplementary Figure S6. Peptide-spectrum matches of MaxQuant and DeepNovo for 3 candidate neoantigens that are likely to be false positives.

Fraction: 20141208_QEp7_MiBa_SA_HLA-I-p_MM15_4_B.raw
Scan ID: 49534
Retention time: 82.974
M/z: 564.327
Charge: 2

### MaxQuant

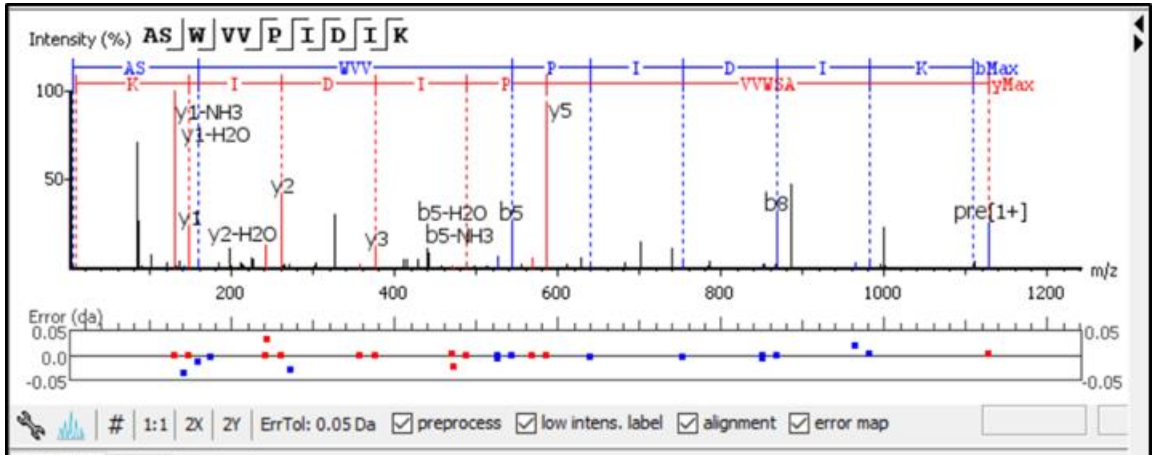


| # | b | b-H2O | b-NH3 | b (2+) | Seq | y | y-H2O | y-NH3 | y (2+) | # |
|---|---|---|---|---|---|---|---|---|---|---|
| 1 | 72.04 | 54.03 | 55.02 | 36.52 | A | | | | | 10 |
| 2 | 159.09 | 141.10 | 142.05 | 80.04 | S | 1056.61 | 1038.60 | 1039.58 | 528.80 | 9 |
| 3 | 345.16 | 327.15 | 328.13 | 173.08 | W | 969.58 | 951.57 | 952.55 | 485.29 | 8 |
| 4 | 444.22 | 426.21 | 427.20 | 222.61 | V | 783.50 | 765.49 | 766.47 | 392.25 | 7 |
| 5 | 543.30 | 525.29 | 526.27 | 272.18 | V | 684.43 | 666.42 | 667.40 | 342.71 | 6 |
| 6 | 640.35 | 622.34 | 623.32 | 320.67 | P | 585.36 | 567.35 | 568.33 | 293.18 | 5 |
| 7 | 753.44 | 735.42 | 736.40 | 377.21 | I | 488.31 | 470.29 | 471.30 | 244.65 | 4 |
| 8 | 868.46 | 850.44 | 851.44 | 434.73 | D | 375.22 | 357.21 | 358.20 | 188.11 | 3 |
| 9 | 981.54 | 963.51 | 964.51 | 491.27 | I | 260.20 | 242.19 | 243.14 | 130.60 | 2 |
| 10 | | | | | K | 147.11 | 129.10 | 130.09 | 74.06 | 1 |

### DeepNovo

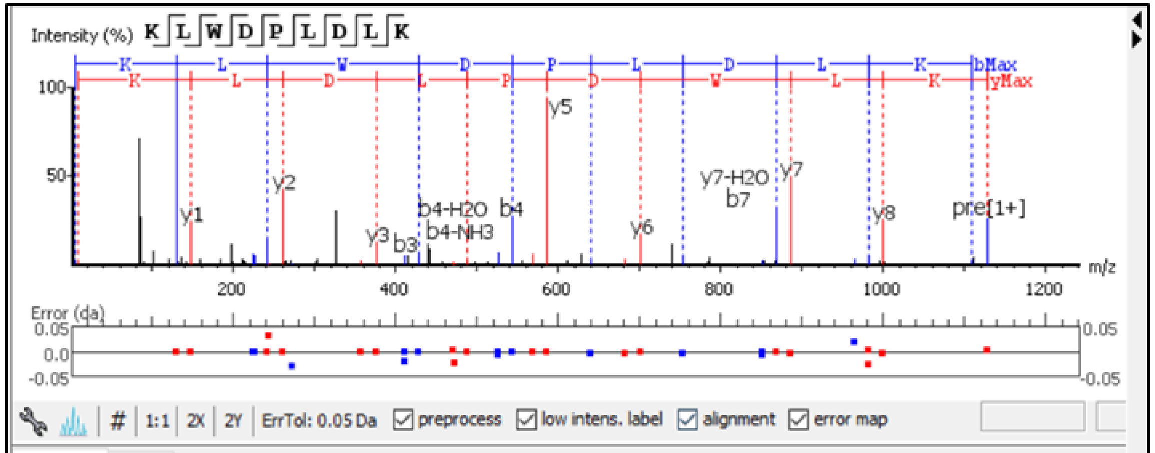


| # | b | b-H2O | b-NH3 | b (2+) | Seq | y | y-H2O | y-NH3 | y (2+) | # |
|---|---|---|---|---|---|---|---|---|---|---|
| 1 | 129.10 | 111.09 | 112.08 | 65.05 | K | | | | | 9 |
| 2 | 242.19 | 224.18 | 225.16 | 121.59 | L | 999.56 | 981.54 | 982.55 | 500.28 | 8 |
| 3 | 428.27 | 410.26 | 411.26 | 214.63 | W | 886.47 | 868.46 | 869.44 | 443.73 | 7 |
| 4 | 543.30 | 525.29 | 526.27 | 272.18 | D | 700.39 | 682.38 | 683.36 | 350.69 | 6 |
| 5 | 640.35 | 622.34 | 623.32 | 320.67 | P | 585.36 | 567.35 | 568.33 | 293.18 | 5 |
| 6 | 753.44 | 735.42 | 736.40 | 377.21 | L | 488.31 | 470.29 | 471.30 | 244.65 | 4 |
| 7 | 868.46 | 850.44 | 851.44 | 434.73 | D | 375.22 | 357.21 | 358.20 | 188.11 | 3 |
| 8 | 981.54 | 963.51 | 964.51 | 491.27 | L | 260.20 | 242.19 | 243.14 | 130.60 | 2 |
| 9 | | | | | K | 147.11 | 129.10 | 130.09 | 74.06 | 1 |

Fraction: 20141210_QEp7_MiBa_SA_HLA-I-p_MM15_2_B_1.raw
Scan ID: 21931
Retention time: 37.371
M/z: 331.854
Charge: 3

### MaxQuant

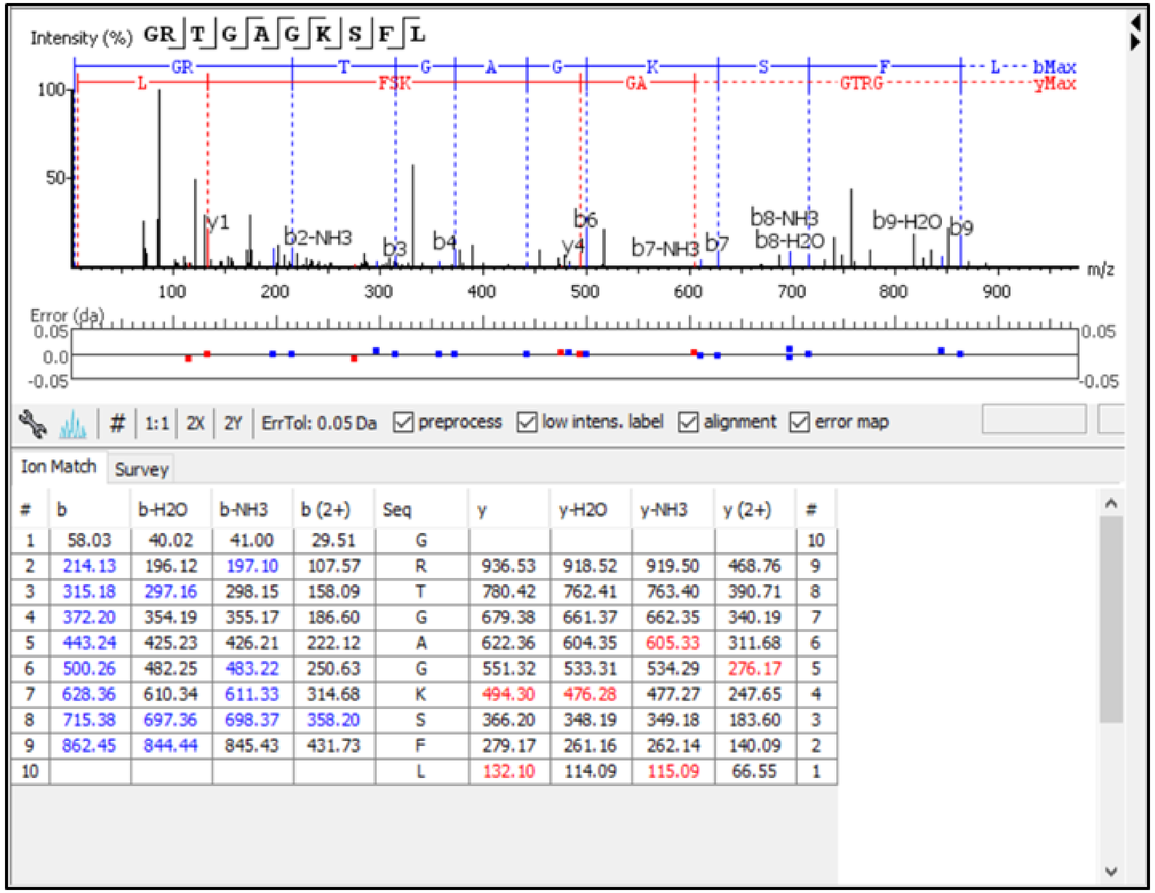


| # | b | b-H2O | b-NH3 | b (2+) | Seq | y | y-H2O | y-NH3 | y (2+) | # |
|---|---|---|---|---|---|---|---|---|---|---|
| 1 | 58.03 | 40.02 | 41.00 | 29.51 | G | | | | | 10 |
| 2 | 214.13 | 196.12 | 197.10 | 107.57 | R | 936.53 | 918.52 | 919.50 | 468.76 | 9 |
| 3 | 315.18 | 297.16 | 298.15 | 158.09 | T | 780.42 | 762.41 | 763.40 | 390.71 | 8 |
| 4 | 372.20 | 354.19 | 355.17 | 186.60 | G | 679.38 | 661.37 | 662.35 | 340.19 | 7 |
| 5 | 443.24 | 425.23 | 426.21 | 222.12 | A | 622.36 | 604.35 | 605.33 | 311.68 | 6 |
| 6 | 500.26 | 482.25 | 483.22 | 250.63 | G | 551.32 | 533.31 | 534.29 | 276.17 | 5 |
| 7 | 628.36 | 610.34 | 611.33 | 314.68 | K | 494.30 | 476.28 | 477.27 | 247.65 | 4 |
| 8 | 715.38 | 697.36 | 698.37 | 358.20 | S | 366.20 | 348.19 | 349.18 | 183.60 | 3 |
| 9 | 862.45 | 844.44 | 845.43 | 431.73 | F | 279.17 | 261.16 | 262.14 | 140.09 | 2 |
| 10 | | | | | L | 132.10 | 114.09 | 115.09 | 66.55 | 1 |

### DeepNovo & PEAKS DB

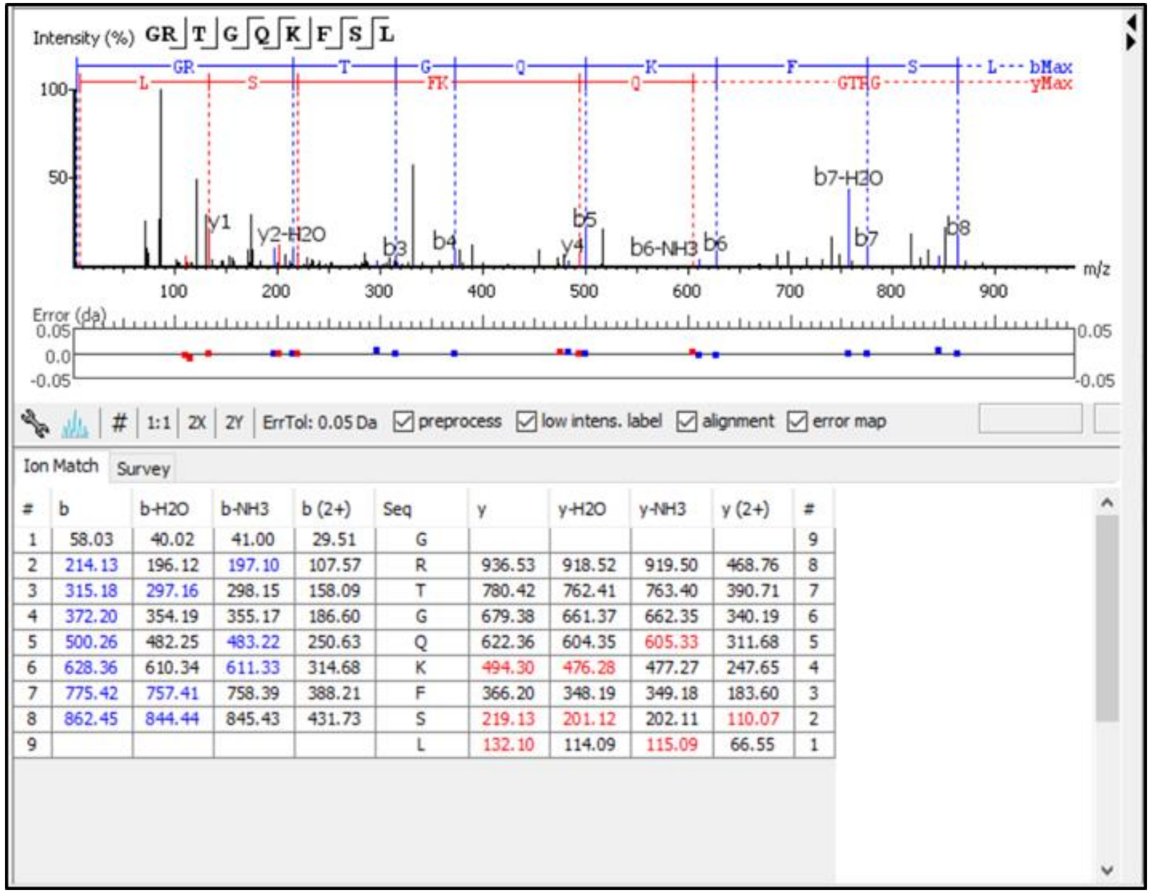


| # | b | b-H2O | b-NH3 | b (2+) | Seq | y | y-H2O | y-NH3 | y (2+) | # |
|---|---|---|---|---|---|---|---|---|---|---|
| 1 | 58.03 | 40.02 | 41.00 | 29.51 | G | | | | | 9 |
| 2 | 214.13 | 196.12 | 197.10 | 107.57 | R | 936.53 | 918.52 | 919.50 | 468.76 | 8 |
| 3 | 315.18 | 297.16 | 298.15 | 158.09 | T | 780.42 | 762.41 | 763.40 | 390.71 | 7 |
| 4 | 372.20 | 354.19 | 355.17 | 186.60 | G | 679.38 | 661.37 | 662.35 | 340.19 | 6 |
| 5 | 500.26 | 482.25 | 483.22 | 250.63 | Q | 622.36 | 604.35 | 605.33 | 311.68 | 5 |
| 6 | 628.36 | 610.34 | 611.33 | 314.68 | K | 494.30 | 476.28 | 477.27 | 247.65 | 4 |
| 7 | 775.42 | 757.41 | 758.39 | 388.21 | F | 366.20 | 348.19 | 349.18 | 183.60 | 3 |
| 8 | 862.45 | 844.44 | 845.43 | 431.73 | S | 219.13 | 201.12 | 202.11 | 110.07 | 2 |
| 9 | | | | | L | 132.10 | 114.09 | 115.09 | 66.55 | 1 |

Fraction: 20141208_QEp7_MiBa_SA_HLA-I-p_MM15_3_B.raw
Scan ID: 2606
Retention time: 6.317
M/z: 334.217
Charge: 3

### MaxQuant

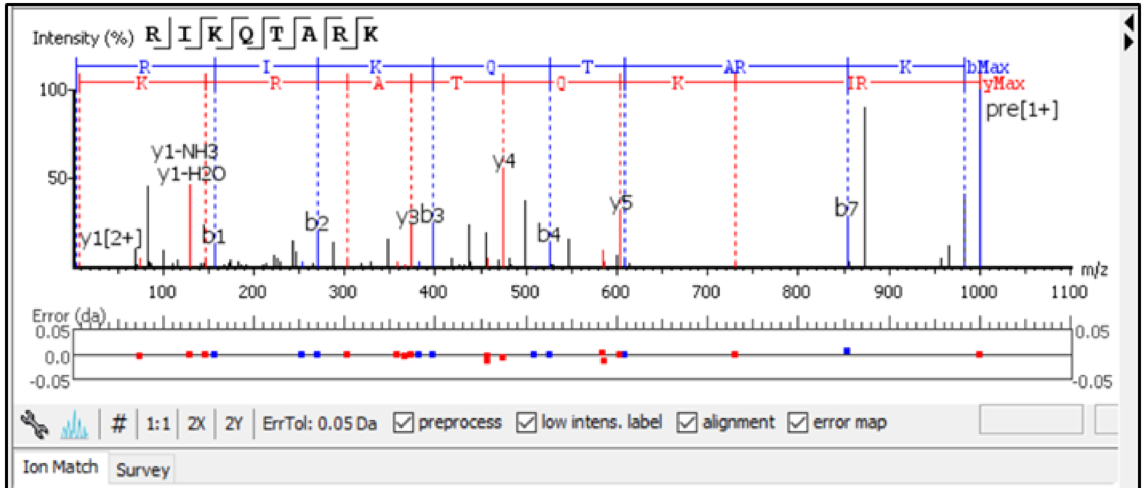


| # | b | b-H2O | b-NH3 | b (2+) | Seq | y | y-H2O | y-NH3 | y (2+) | # |
|---|---|---|---|---|---|---|---|---|---|---|
| 1 | 157.11 | 139.10 | 140.08 | 79.05 | R | | | | | 8 |
| 2 | 270.19 | 252.18 | 253.17 | 135.60 | I | 844.54 | 826.53 | 827.51 | 422.77 | 7 |
| 3 | 398.29 | 380.28 | 381.26 | 199.64 | K | 731.45 | 713.44 | 714.43 | 366.23 | 6 |
| 4 | 526.35 | 508.34 | 509.32 | 263.67 | Q | 603.36 | 585.34 | 586.34 | 302.18 | 5 |
| 5 | 627.39 | 609.38 | 610.37 | 314.20 | T | 475.31 | 457.30 | 458.28 | 238.15 | 4 |
| 6 | 698.43 | 680.42 | 681.40 | 349.72 | A | 374.25 | 356.24 | 357.23 | 187.63 | 3 |
| 7 | 854.52 | 836.52 | 837.51 | 427.77 | R | 303.22 | 285.20 | 286.19 | 152.11 | 2 |
| 8 | | | | | K | 147.11 | 129.10 | 130.09 | 74.06 | 1 |

### DeepNovo

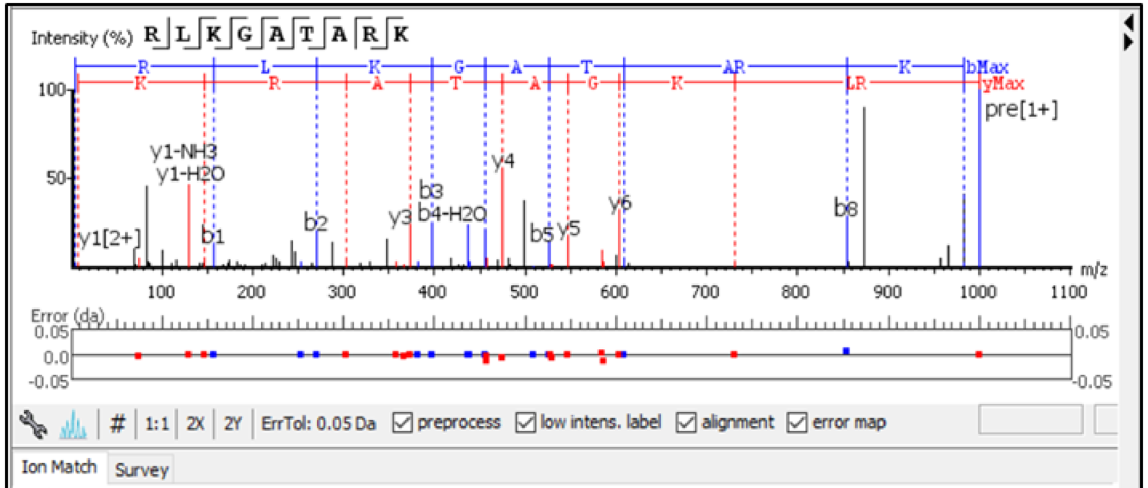


| # | b | b-H2O | b-NH3 | b (2+) | Seq | y | y-H2O | y-NH3 | y (2+) | # |
|---|---|---|---|---|---|---|---|---|---|---|
| 1 | 157.11 | 139.10 | 140.08 | 79.05 | R | | | | | 9 |
| 2 | 270.19 | 252.18 | 253.17 | 135.60 | L | 844.54 | 826.53 | 827.51 | 422.77 | 8 |
| 3 | 398.29 | 380.28 | 381.26 | 199.64 | K | 731.45 | 713.44 | 714.43 | 366.23 | 7 |
| 4 | 455.31 | 437.30 | 438.28 | 228.15 | G | 603.36 | 585.34 | 586.34 | 302.18 | 6 |
| 5 | 526.35 | 508.34 | 509.32 | 263.67 | A | 546.34 | 528.33 | 529.32 | 273.67 | 5 |
| 6 | 627.39 | 609.38 | 610.37 | 314.20 | T | 475.31 | 457.30 | 458.28 | 238.15 | 4 |
| 7 | 698.43 | 680.42 | 681.40 | 349.72 | A | 374.25 | 356.24 | 357.23 | 187.63 | 3 |
| 8 | 854.52 | 836.52 | 837.51 | 427.77 | R | 303.22 | 285.20 | 286.19 | 152.11 | 2 |
| 9 | | | | | K | 147.11 | 129.10 | 130.09 | 74.06 | 1 |
