## Supplementary Figures and Tables for "Personalized deep learning of individual immunopeptidomes to identify neoantigens for cancer vaccines": Table_S1.pdf

Table S1. Personalized de novo sequencing workflow of neoantigen discovery for patient Mel-15.

| Details of each step in the workflow | HLA-I | HLA-II |
|---|---|---|
| Step 1: Build the immunopeptidome of the patient | | |
| Number of identified peptide-spectrum matches | 341,216 | 67,021 |
| Number of identified database peptides | 35,551 | 9,664 |
| Number of unlabeled spectra | 596,915 | 135,490 |
| Step 2: Train personalized machine learning model | | |
| Number of training PSMs | 307,058 | 60,822 |
| Number of validation PSMs | 17,217 | 2,999 |
| Number of test PSMs | 16,941 | 3,200 |
| Step 3: Personalized de novo sequencing | | |
| Number of raw de novo peptides | 441,274 | 93,983 |
| Step 4: Quality control | | |
| Number of high-confidence de novo peptides | 16,226 | 2,717 |
| Number of de novo peptides at 1% FDR | 5,320 | 863 |
| Step 5: Neoantigen selection | | |
| Missense mutations with at least 2 PSMs | 177 | 70 |

PSM: Peptide-Spectrum Match

FDR: False Discovery Rate
