## Supplementary figures and images for "Personalized deep learning of individual immunopeptidomes to identify neoantigens for cancer vaccines"

### Figure_S4.pdf

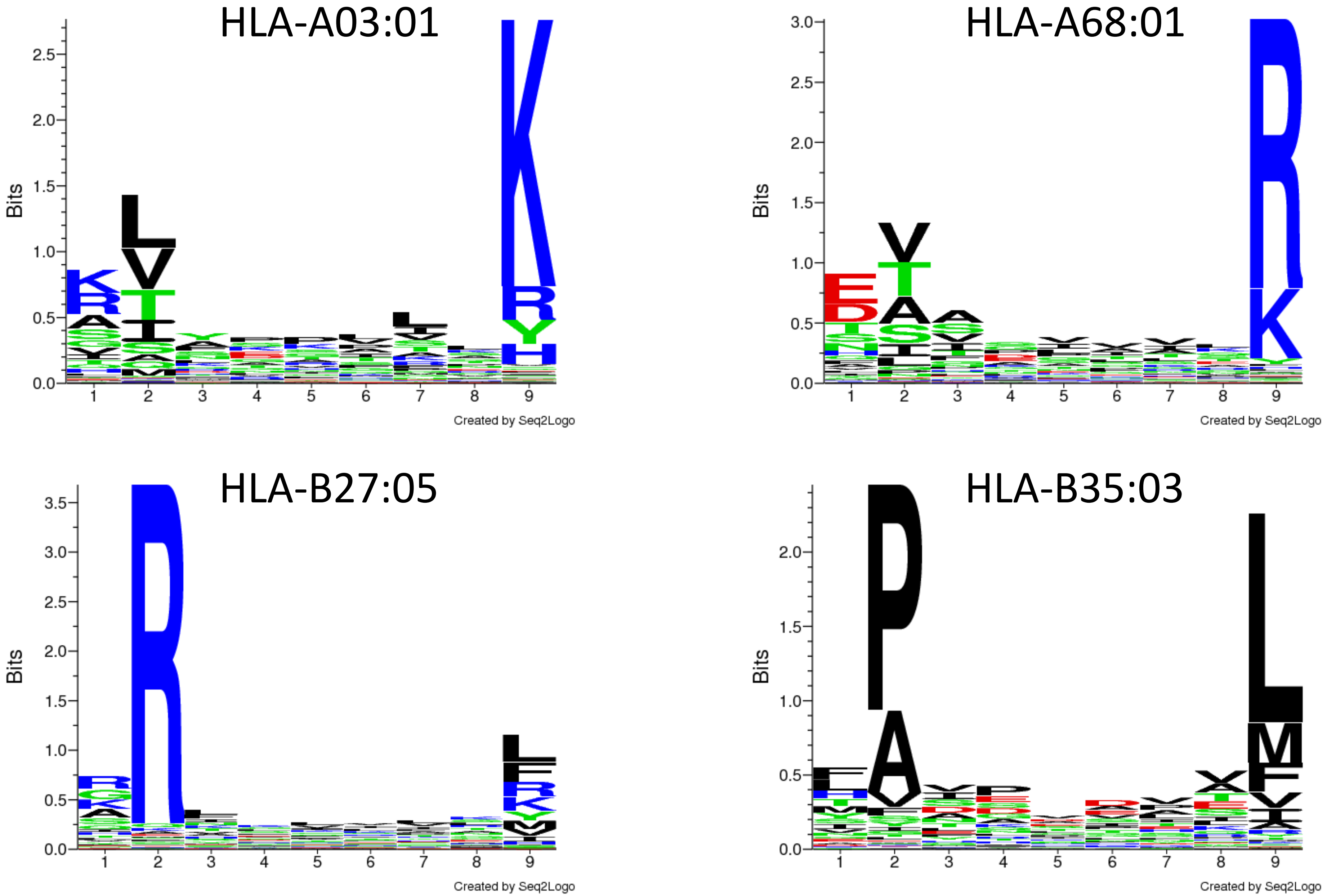


Supplementary Figure S4. Binding motifs of database HLA-I peptides of patient Mel-15.

### Figure_S5.pdf

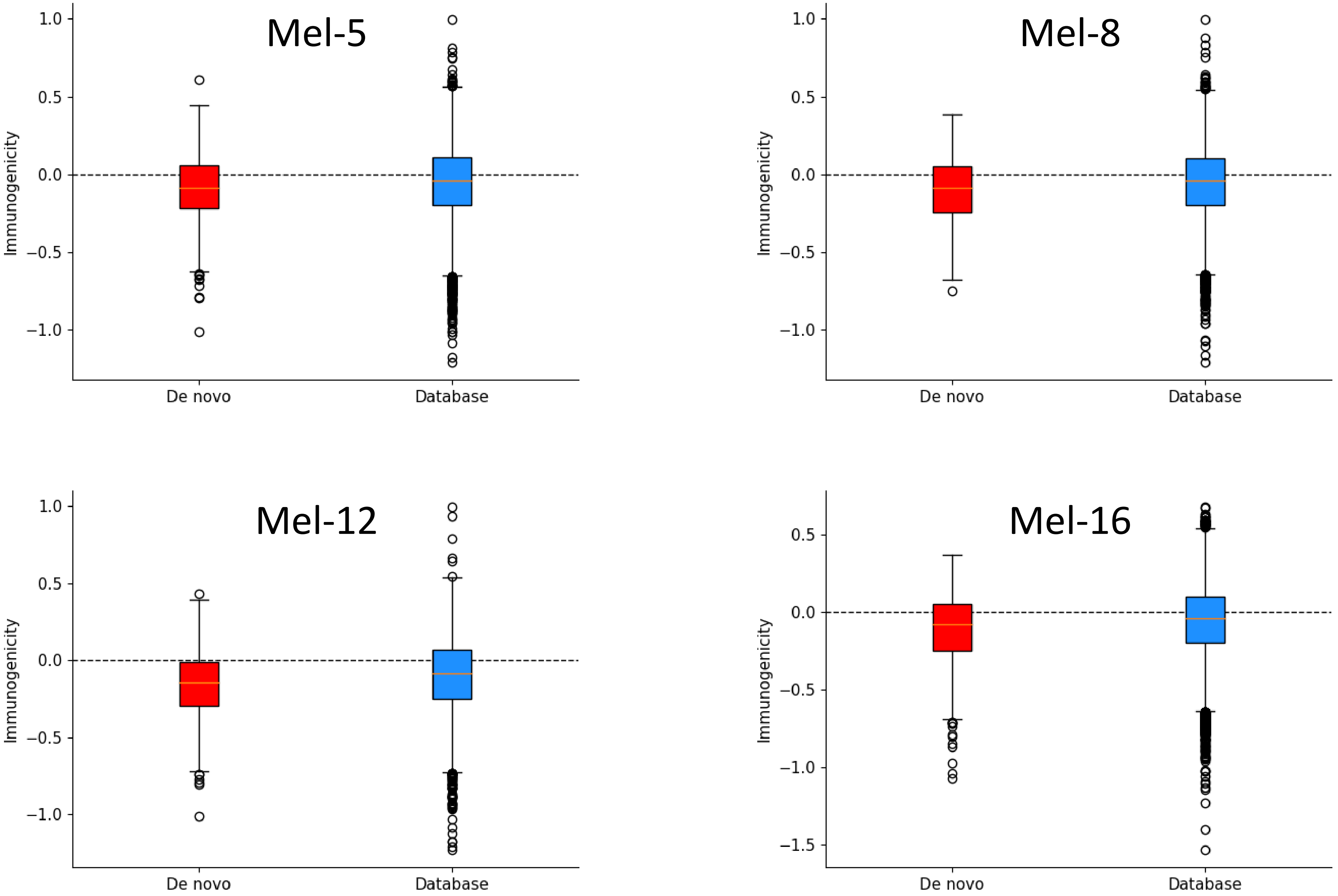


Supplementary Figure S5. Immunogenicity of *de novo* and database HLA-I peptides.
